# The adenomatous polyposis coli protein 3o years on

**DOI:** 10.1101/2022.11.14.516391

**Authors:** James Abbott, Inke S. Näthke

## Abstract

Mutations in the gene encoding the Adenomatous polyposis coli protein (APC) were discovered as driver mutations in colorectal cancers almost 30 years ago. Since then, the importance of APC in normal tissue homeostasis has been confirmed in a plethora of other (model) organisms spanning a large evolutionary space. APC is a multifunctional protein, with roles as a key scaffold protein in complexes involved in diverse signalling pathways, most prominently the Wnt signalling pathway. APC is also a cytoskeletal regulator with direct and indirect links to and impacts on all three major cytoskeletal networks. Here, we interrogate the enormous depth of sequencing data now available to reveal the conservation of APC across taxonomy and relationships between different APC protein families.

---

Mutations in the gene encoding the Adenomatous polyposis coli protein (APC) were discovered as driver mutations in colorectal cancers almost 30 years ago (1–4) initially in mouse models and also patients (5). Since then, the importance of APC in normal tissue homeostasis has been confirmed in a plethora of other (model) organisms including c. elegans, Drosophila, rats, zebra fish, pigs, and more (6–9). A key role for APC is its ability to act as a key scaffold protein in complexes involved in diverse signalling pathways. Most prominent is the Wnt signalling pathway. Here, APC is a crucial player in assembling the protein complex that phosphorylates b-catenin targeting it for degradation in the absence of Wnt signals. In the presence of Wnt signals, this complex is inactivated, b-catenin can accumulate and direct transcriptional changes that are associated with proliferative, less differentiated states (10). In addition, APC is a cytoskeletal regulator with direct and indirect links to and impacts on all three major cytoskeletal networks (11).

Both the function of APC in Wnt signalling and in cytoskeletal regulation and the resulting contribution to the behaviour of cells and tissues have been extensively summarised and reviewed and highlight the complexity of the interactions of the APC protein and their outputs for cellular function (11). Studies interrogating the APC interactions directly, further support the diversity and complexity of its links to many different cellular processes (12). More recently, the idea that APC itself can undergo phase separation adds another layer of complexity to its regulation and could be a means for APC to response to local intracellular conditions as has been shown in other disordered proteins (13; 14). It is thus not surprising that mutations in this one gene can have such profound effects, particularly on the lining of the intestinal tract, the most dynamic tissue in the body.

Additional complexity is introduced by the fact that there two related but distinct APC proteins, APC and APC2 have been described (15). The relationship between them is not entirely clear. Mutations in cancers, particularly colorectal cancers, are extremely common only in the former although both seem able to support Wnt signalling and there have been some reports about APC2 also able to affect the cytoskeleton (16–18). The majority of research has focussed on APC and there is comparably little information available about APC2. One exception is in Drosophila where both of the expressed APC proteins have been investigated. However, how each relate to APC and APC2 is not entirely clear cut (see below).

The human intestinal tract is estimated to shed 20-50 million cells per minute leading to the renewal of the entire lining every five days (19). Normal homeostasis relies on stem cells that produce transit amplifying cells to generate the complement of cell types that constitute the intestinal epithelium. Most prominent are absorptive enterocytes, that are protected by mucus-secreting Goblet cells, enteroendocrine and Paneth cells. The latter are found in intestinal crypts, where they provide crucial factors to create the stem cell niche environment, including Wnt and Notch (20; 21). Similar cells are also located in colonic crypts (22). Constant production of new cells from stem cells, differentiation into different lineages, and shedding from the tissue layer have to be well balanced for normal tissue function. In addition, the system can respond to injury and rapidly replace damaged areas to maintain the crucial barrier function that prevents entry of pathogens from the lumen of the intestinal tract. Directed migration of cells from crypts towards the lumen is an integral feature of these processes (23). Directed migration is also important for the rapid closure of any areas denuded of the epithelial layer in response to injury and inflammation (24). Integrating key signalling pathways allows careful tuning of this system to create required responses in a locally and temporally regulate manner. The ability of APC to interact with many different proteins and thus contribute to and coordinate many different signalling pathways, makes it a key integrator of such external signals. It plays an important role in coordinating the cellular responses required for normal homeostasis.

Mutations in APC in cancer most commonly produce truncated APC protein of varying lengths. Such truncated APC proteins lack many of its interaction sites, reducing its ability to integrate incoming signals. It is thus not surprising that loss of APC has profound effects on intestinal epithelial organisation, including loss of differentiation and changes in tissue shape (25–27) to initiate tumours. Losing fully functional APC impacts on all the processes required for normal homeostasis of a rapidly renewing epithelium: it activates Wnt signalling – promoting proliferation at the expense of differentiation, it reduces mitotic fidelity – increasing genetic instability, compromises directed cell migration – increasing the residence time of APC-mutant cells in crypts, and it changes the direction of detachment of cells from the basal layer – reducing sloughing off into the gut lumen. The latter two consequences directly bestow a competitive advantage over wild type cells by increasing the probability of APC-mutant cells to remain in the tissue while wild type cells are removed.

Unsurprisingly given the many processes APC impacts, many binding partners have been identified. The APC protein in humans contains 2,843 amino acids (some alternatively spliced forms have been reported but here we will focus on the most commonly described form). Many APC interactions and their regulation have been described individually (11; 28) (Figure 1 A) and binding sites for many of its binding partners have been mapped (28). Those relating to Wnt signalling and some microtubule interactions are summarised in Figure 1 A. The complexity of APC interactions is consistent with the idea that the context of individual interactions affect each other (29). In other words, there are likely many combinations of interactions that can occur simultaneously. These will depend on the subcellular and molecular conditions and context but relatively little is known about the combinations that do and do not occur. Two thirds of the APC protein is easily degraded and was predicted to be disordered by early biochemical experiments and resulting models (30). Much has been learned about disordered protein domains in the last years and the tools to analyse them have become increasingly sophisticated. Intrinsically disordered domains, including some in APC, have been proposed to be highly dynamic and to provide interchangeable interaction sites for different partners to create a “cloud of bound conformations” (31). Intrinsically disordered regions have also been suggested to act as localised sensors to different intracellular conditions (14; 32). One response can be for such domains to forming liquid phase separated structures (14; 32). This adds an additional layer of regulation of the functions APC can perform.

**Figure 1:**
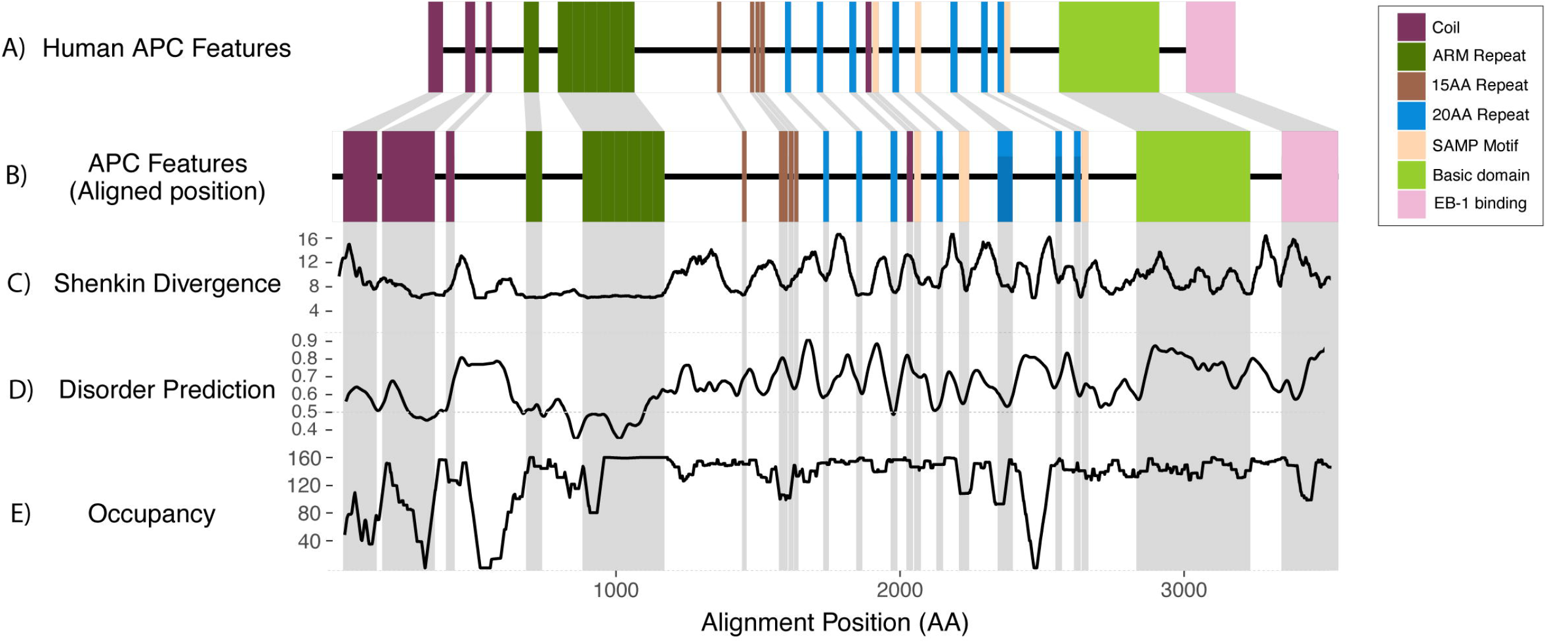
APC features in the context of multiple sequence alignment of 158 Chordata orthologs. A. Simplified APC domain structure, showing domains involved in Wnt signaling and microtubule binding. Co-ordinates are absolute co-ordinates of the Human APC sequence (P25054) B. Domain structure from A), with co-ordinates projected onto location in multiple sequence alignment C. Windowed Shenkin diversity scores, determined using a 50 amino acid rolling window. The Shenkin divergence score gives an indication of how divergent the sequences are at each position in the alignment, within a range of 4-106. D. Windowed disorder prediction determined using JRONN with a 50 amino acid rolling window. Values above 0.5 indicate a potentially disordered region, with higher scores indicating an increased probability of the region being disordered. E. Windowed occupancy determined using a 50 amino acid rolling window. This represents the number of sequences at each position within the alignment, which do not contain a gap.

Currently, how APC’s diverse interactions are coordinated and regulated remains largely unknown. Similarly, how disease-associated mutations in APC affect the balance of all the responses it can integrate remains under investigated. The prevalence of mutations in the APC gene places this question at the heart of understanding this important player in cell behaviour. Answering this question will help to develop full and accurate predictions of the effectiveness of potential therapeutic or prognostic tools that aim to target such mutations. Using the molecular information available to identify common and different themes in the diverse organism where APC has been identified and studied is an important first step. To that end we interrogated and compared sequence of currently known APC proteins.

OrthoDB (https://www.ezlab.org/orthodb.html; (33) provides curated sets of predicted orthologs from a wide variety of organisms, determined through a best-reciprocal blast and clustering approach. InterPro (www.ebi.ac.uk/interpro; (34) produces protein signatures using a range of databases of protein domains and sites, enabling families of proteins to be identified based upon the presence of a number of representatives of a signature. Mappings of InterPro signatures to members of the UniProt database (35) provides an alternative approach to identifying related proteins. A single InterPro family <u>(IPR026818)</u> represents all APC sequences spanning a broad range of taxonomy from primitive phyla such as Cnidaria and Porifera throughout the Chordata, including Aves (birds), Mammalia and Actinopteri (ray-finned fish). The family incorporates two subfamilies containing representatives of APC <u>(IPR026836)</u>, and APC2 <u>(IPR026837)</u>, but also includes sequences that do not fall into either of these families, including APC-related protein 1 (APR1) expressed in C. Elegans and the two proteins expressed in Drosophila (Figure 2).

**Figure 2:**
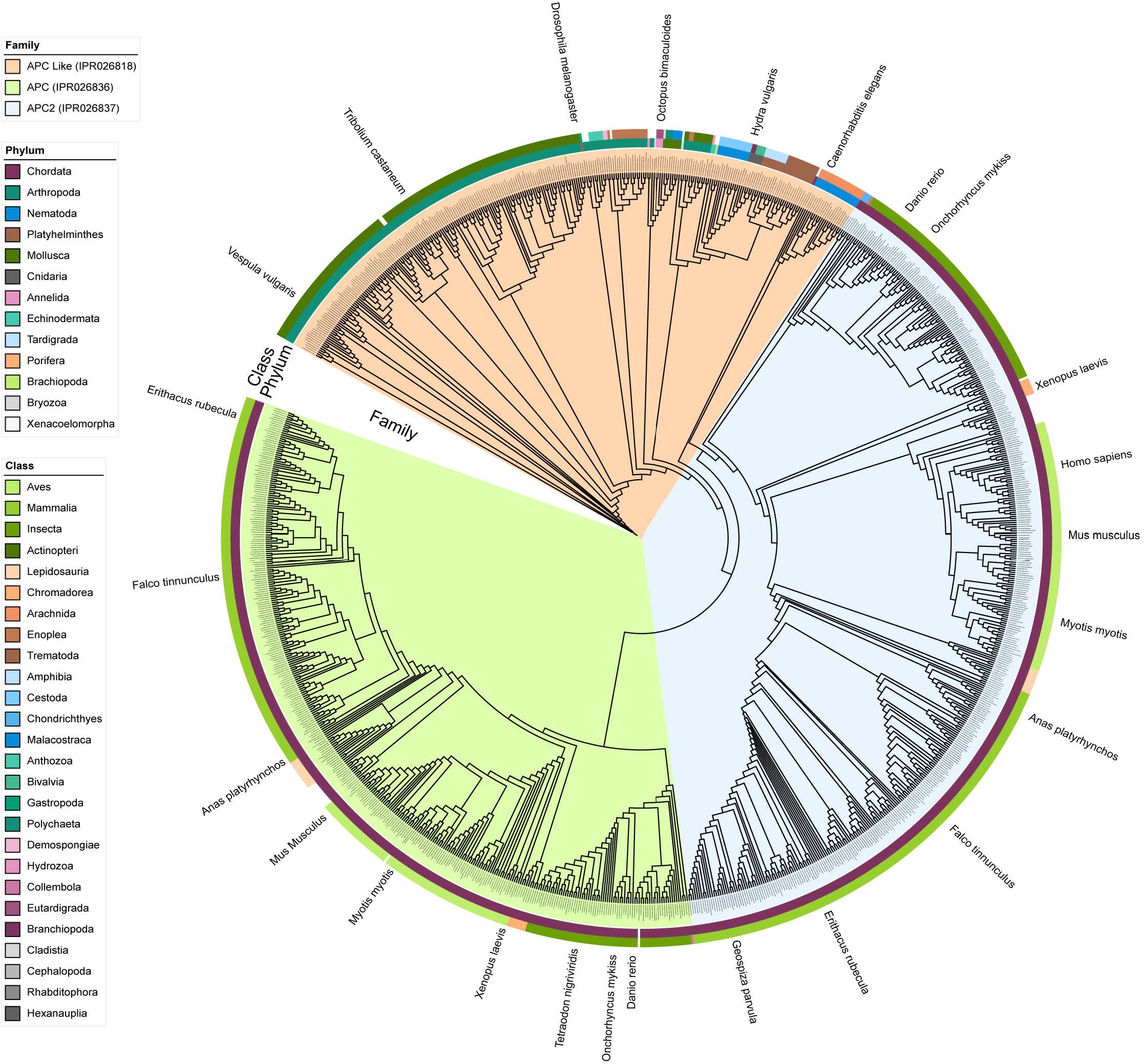
Taxonomy of APC protein family members. The colour of the clades indicates the APC family to which each sequence belongs. The inner coloured ring indicates the phylum of the organism, and the outer ring the class. Due to the extreme degree of conservation within the APC and APC2 families branch lengths are ignored in this figure to enable the structure to be evident. A separate version which utilises branch lengths is included as supplemental figure 2.

The emergence of new tools to interrogate proteins computationally and our increased ability to predict structure – function relationships provide an opportunity to revisit this protein and its properties, which have remained somewhat enigmatic

To that end, in this review, we aim to collate information currently available about the molecular details of APC across taxonomies, identify robust structural features, and compare and contrast APC and APC2 to increase our understanding of its properties.

## METHODS

To gain an indication of the conservation of APC, a set of metazoan APC orthologs were downloaded from OrthoDB 10.1 (33) (entry 21888at33208). These were refined according to their annotations to remove APC2 sequences, which are included in the same ortholog set as APC, those annotated as low quality, and also those which were other than the canonical isoform. The sequences were subsequently filtered to remove sequence shorter than 2750 AA, to remove partial length sequences, and also to remove duplicates from the same organism. Multiple sequence alignment was carried out using ClustalW 2.1 (36)with default parameters. The alignment was then edited using Jalview 2.11.2.4 (37)and outlying sequences, based upon a principal component analysis, removed. Shenkin diversity scores were determined using AAcon 1.1 (38) and disorder predictions carried out using JRonn ((39; 40)) within Jalview. Features annotated in the Uniprot record for the Human APC sequence (P25054) and InterPro domain mappings were projected onto the alignment positions along with the Shenkin diversity and JRonn disorder predictions (Figure 1C and 1D).

All sequences belonging to the InterPro Adenomatous polyposis coli (APC) family (IPR026818), which includes APC, APC2 and APC-like sequences were downloaded from InterPro release 89.0 (34). Those sequences belonging to the APC family (IPR026836) were filtered to remove partial length sequences shorter than 2,000AA, while sequences in the APC2 family (IPR026837) were similarly filtered with a 1,000AA cut-off. Redundancy was removed from the group by removing identical sequences from the same organisms. The sequence set includes multiple isoforms, and sometimes duplicated sequences from different assemblies or sequencing projects, so the sequences were refined using a custom Python script. The UniProt record for each sequence was parsed to identify records in DNA sequence databases from which the protein sequence was derived, and both the genomic locus and assembly version identified. In cases where multiple assemblies were present, sequences from a single assembly were retained, with assemblies present in the Ensembl (41) database preferred, as was the longest protein sequence associated with each genomic locus. In addition to removing redundancy from the dataset, this process also allowed the identification of instances where genes were duplicated within an assembly, since the remaining protein sequences may be associated with multiple genomic loci within the same assembly. This resulted in a set of 1253 sequences covering 910 species.

A multiple sequence alignment was carried out as previously described, then phylogenetically informative regions were selected from the alignment using BMGE (42) with a Blosum62 similarity matrix, then a phylogenetic tree was then constructed using RaxML-NG 1.1 (43)using a JTT+G model with 1,000 bootstrap iterations, an MRE-based bootstrap convergence metric and Felsenstein branch support metric. The resulting tree was visualised and annotated using the Interactive Tree of Life (44).

The three-dimensional structures of the Alphafill (45) models for Human APC and APC2 were visualised using Jalview and ChimeraX (46), with the intrinsically disordered regions downstream of the Armadillo repeats (residues beyond 800AA) hidden for clarity.

## RESULTS

- The multiple sequence alignment of 158 Chordata APC sequences shows an extremely high degree of conservation. The distribution of Shenkin divergence scores along the length of the alignment shows that the annotated domains coincide with more conserved regions (Figure 1 B/C). These domains also tend to occur in regions with lower predicted disorder (Figure 1 D), even in the highly disordered region associated with β-catenin binding. There appears a clear association between these known binding domains and increased sequence conservation, and decreased disorder.
- Phylogenetic analysis shows a clear separation between the three InterPro families and their members (Figure 2; Interactive versions of trees available online at https://itol.embl.de/shared/jca): APC, APC2, APC-like. APC and APC2 are found exclusively in the Chordata, while APC-like sequences are found only in the ‘lower animals’, including Insecta, Amphibia, Hydrazao and Trematoda.
- Members of the APC family (IPR026836) show extensive conservation across the length of the protein, including disordered regions (Mean Shenkin Divergence score: 8.66, SD 4.24) (For reference: complete conservation would be indicated by a score of 4) (Figures 1 and 2). Such extreme conservation across the entire sequence of APC suggests that all residues and their relative position to each other is crucial for the full complement of its functions and their coordination. This is consistent with the importance for APC to interact with many different binding partners.
- The APC2 family is also highly conserved, but to a lesser degree than APC (Mean Shenkin Divergence score: 11.25, SD 6.65) (Figures 2).
- The proteins in the APC-like family show far greater divergence than those in the APC or APC2 families (Figure 2).
- The vast majority of organisms have a single copy of the APC and APC2 genes (Supplemental Figure 1). In a small number of species, two copies of the genes were observed, primarily in the teleost fish i.e. rainbow trout, Atlantic salmon (supplementary Table 1, and supplementary Figure 3), although there are also examples of duplications within Aves and Amphibia. The duplications within the teleost fish are likely a consequence of a genome duplication, which occurred ~90 million years ago, resulting in the ancestral species carrying a tetraploid genome, which is believed to be going through a process of diploidisation (47). The second copy of APC within these species form separate sub-groups within the phylogeny (Supplementary Figure 1), which have independently diverged following the whole genome duplication, while still retaining a high degree of conservation. Duplicated APC2 sequences, show a similar pattern of duplication. The majority of species showing duplications have both APC and APC2 duplicated, although some species carry duplicates of only one (Supplementary Figure 2).
- The APC and APC2 families are only found in Chordata and are broadly distributed across the phylum. All other phyla that have APC fall into the APC-like group, where diversity is considerably higher (Figure 2).
- Amongst the Chordata, which have either APC, APC2 or both, but never an APC-Like gene, 62.1 % of organisms carry both APC and APC2, while 25% have only APC, and 12.9% have only APC2. (Supplementary Figure 1).
- Alphafill models are available for both Human APC (first 1,400AA) and APC2 (full length model). These are high-confidence models for N-terminal coils and the armadillo repeats. Although the length of the disordered regions is included in the model for APC2, these are only partially represented in the APC model. These disordered regions unsurprisingly are regions with low confidence in the predicted models, and do not have any evident predicted structure. Disordered regions have been proposed to be highly dynamic and to provide interchangeable interaction sites for different partners by creating a “cloud of bound conformations” (31) (Figure 3). One response of changes in intrinsically disordered regions (IDR) can be formation of liquid phase separated structures (14; 32). Indeed, APC has been suggested to do just that (13; 31).
- Uniprot records for the Human APC (P25054) and APC2 (O95996) sequences are annotated with 7 and 6 armadillo repeats respectively. A comparison of the alphafill structures indicates that this region is extremely similar between the two proteins (Figure 3; Supplementary files 1 and 2), with no obvious difference in repeat number. The annotated repeats in both proteins differ; however, eight different repeating units are annotated in total. The difference in repeat number appears to be a result of differing interpretations of what forms an Armadillo repeat by those annotating the sequences, with the main differences in the terminal repeats. It is notable that the annotated repeat units are also different, with the APC2 repeats each starting about halfway through an APC repeat.
- N-terminal coils and armadillo repeats are the most prominent structural features (Figure 3). Using the latter to compare the relative position of structural features in APC and APC2 suggests that the N-terminal coils may be oriented differently relative to armadillo repeats in APC and APC2. Alphafold’s predicted Aligned Error implies that while the N-terminal coils and armadillo repeats are likely correctly predicted, the relative positioning of these is uncertain. Given the established importance of the armadillo repeats for many APC interactions, it is possible that these interactions vary between and are differently regulated in APC2 and APC, and result in different access to the armadillo repeats where many interacting partners bind (Figure 1 and 3).
- Superposition of experimentally determined structures of portions of the APC protein with the Alphafill model using ChimeraX’s MatchMaker tool suggests these structural regions have been predicted extremely well by Alphafold. PDB:5IZA represents the majority of the armadillo repeat region (P25054:407-751), which when superimposed has an RMSD of 0.566 □. Similarly, superposition of three alpha-helices from the annotated N-terminal coils (PDB:1M5I; P25054:126-250) results in an RMSD of 0.896 □.
- The ability of the disordered regions to create flexible clouds around the armadillo repeats could be affected by binding of proteins to the disordered regions. This could explain why truncated forms of APC, lacking much of the disordered regions, are more active. For instance, truncated APC appears to be more active in stimulating ASEF (48) suggesting that a regulatory feature provided by the disordered region is lacking in truncated forms of APC.
- A loop between Arm 6 and 7 in Human APC has two phosphosites (SER744 and SER748) that have been experimentally confirmed (49) (Figure 3). These may be important for positioning the disordered regions beyond the arm repeats. One of these sites is conserved in APC2 (SER710) where it is similarly located in a loop within the final armadillo repeat (Figure 3).

**Figure 3:**
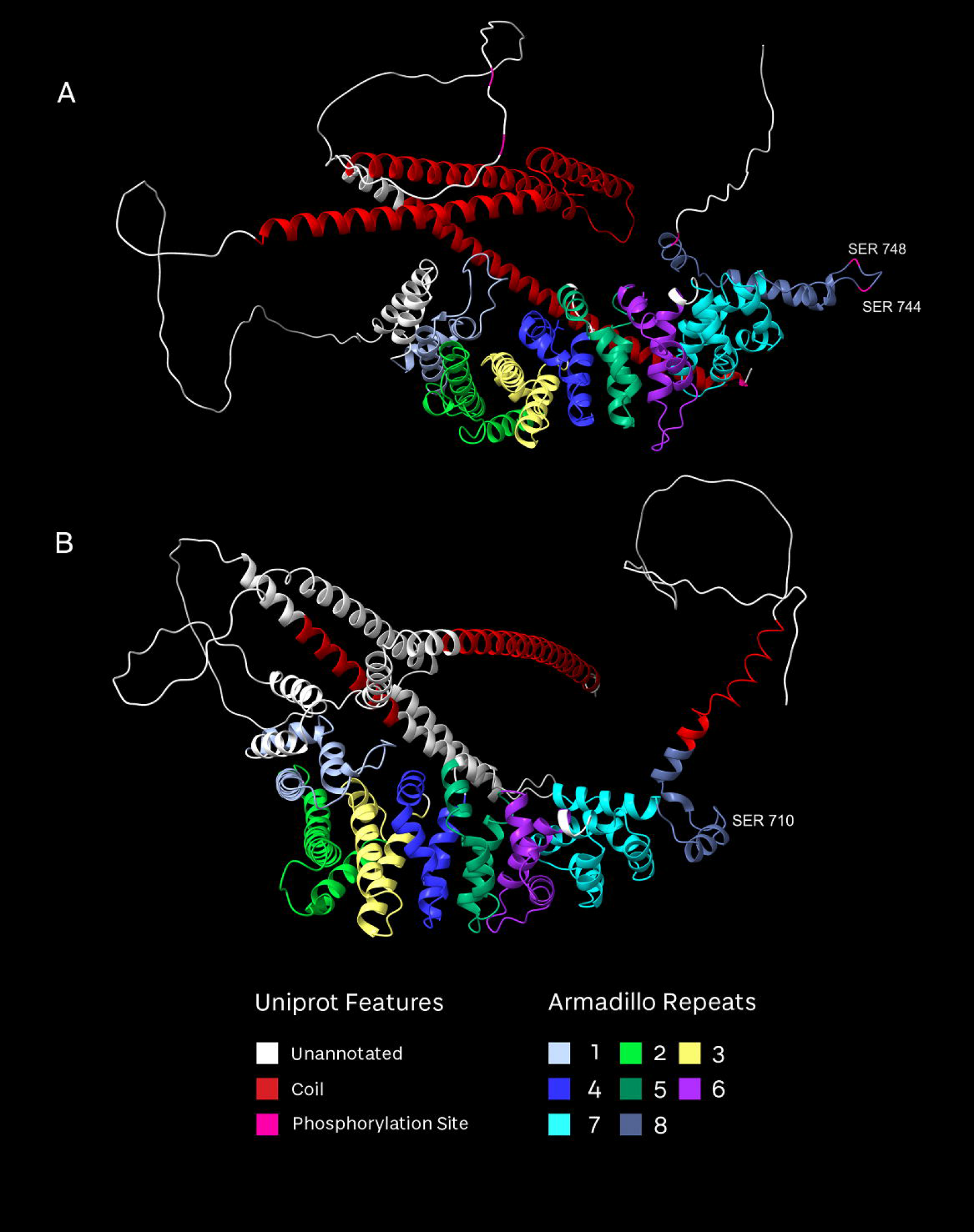
Alphafill structural predictions of the N-terminal region (residues 1-800) of APC (A) and APC2 (B). Helices annotated within the Uniprot records are highlighted in red. The labelled residues are experimentally identified phosphorylated serine residues in APC, and the conserved S744 residue in APC2.

## CONCLUSION

The extreme degree of conservation even across the extensive disordered regions confirms the importance of APC existing as a single protein. Dividing it into smaller proteins that work independently to perform the same functions individually would remove the ability to coordinate them. This is consistent with the idea that the ability to integrate and coordinate many different pathways and processes is a key feature supporting the multifunctionality of APC.

This high conservation across the entire APC sequence also underscores the importance of investigating the complement of APC interactions and potential functions, even when targeting only a single interaction (for instance by deletion mutations). The ability or many of the annotated domains of APC to interact with several different binding partners further illustrates the complex interplay of APC interactions and their regulation. The high content of intrinsically disordered domains provides an extremely dynamic means to create a plethora of conformations to support many different combinations of interactions for APC. Intrinsically disordered domains can respond to different subcellular local conditions to create different conformations, which in turn creates possibilities to regulate individual or combinations of interactions spatially.

The presence of three distinct APC protein families suggests that different organisms have evolved these proteins to be optimised for their needs. Of note is that the APC proteins in Drosophila and C. elegans for instance are members of the APC-like protein family, which may contribute to differences in how they interact with specific binding partners, how they are regulated and the specific details of how they contribute to distinct functions. This also may need to be considered when relating either of the two APC-like proteins in Drosophila to APC and APC2 directly.

Possible differences between the conservation and structure of APC and APC2 suggest there might be differences in their interactions and regulation, particularly in the N-terminal region where the position of coils relative to the armadillo repeats is predicted to vary (Figure 3).

One surprising finding was that the APC-like protein in Hydra vulgaris (Uniprot:T2MGZ0) does not have annotated armadillo repeats; however, the InterPro classifications (https://www.ebi.ac.uk/interpro/protein/UniProt/T2MGZ0/) identify six armadillo repeats at the C-terminus of the protein, whereas these are typically found in close proximity to the N-terminus. Only a single Armadillo ‘repeat’ is annotated in the C.elegans APR-1 sequence, between residues 314-358 of a 1,188AA protein. In neither case are the 15AA/20AA/SAMP domains, which are implicated in binding b-catenin and Axin annotated in the database records. These findings may be a shortcoming of annotation but warrant further investigation.

### The future

We anticipate (and hope) that our analysis and the data provided will help to guide future work to understand and elucidate mechanisms for APC functions. Comparing the conservation of APC binding partners, particularly their APC-binding domains, may shed light on how similar contributions of APC to different signalling pathways are across organisms.

## ACKNOLWEDGEMENTS

We are immensely grateful to S. MacGowan, M. Tsenkov, and G. Barton (Computational Biology, University of Dundee) for immensely helpful discussions.

## CODE AVAILABILITY

The code used in the generation of results and figures is available from https://github.com/bartongroup/30_years_of_APC.

## SUPPLEMENTAL MATERIAL

Supplemental Figure 1: UpSet Plot of distribution of species with APC, APC2, APC-like or combinations of these.

Supplemental Figure 2: Version of Figure 2 that plots branch lengths to indicate degree of conservation evident across the range of taxonomy. Taxonomy of APC protein family members

Supplemental Figure 3: Phylogenetic tree showing APC and APC2 species within the Actinopetri, which include a number of species with duplications of APC and/or APC2. Duplicated proteins are linked with arrows that are colour coded according to the identity of the duplicated sequences.

Supplemental Table 1: Taxonomic details of species included in phylogenetic analysis including copy number of each class of APC, and the Uniprot accessions of the sequences.

