## Supplementary figures and images for "The adenomatous polyposis coli protein 3o years on"

### Supplemental figure 1

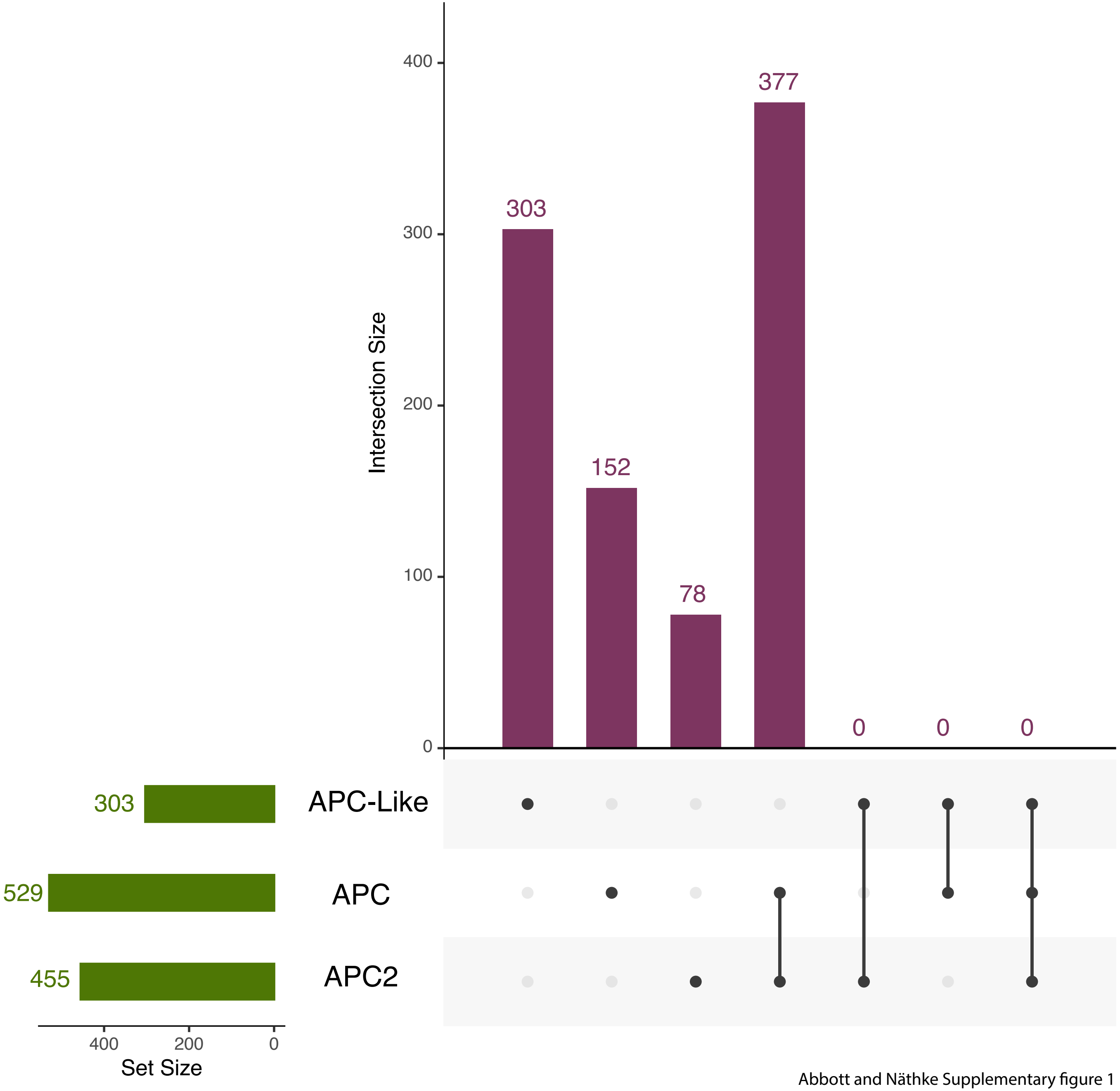


Abbott and Näthke Supplementary figure 1

### Supplemental figure 2

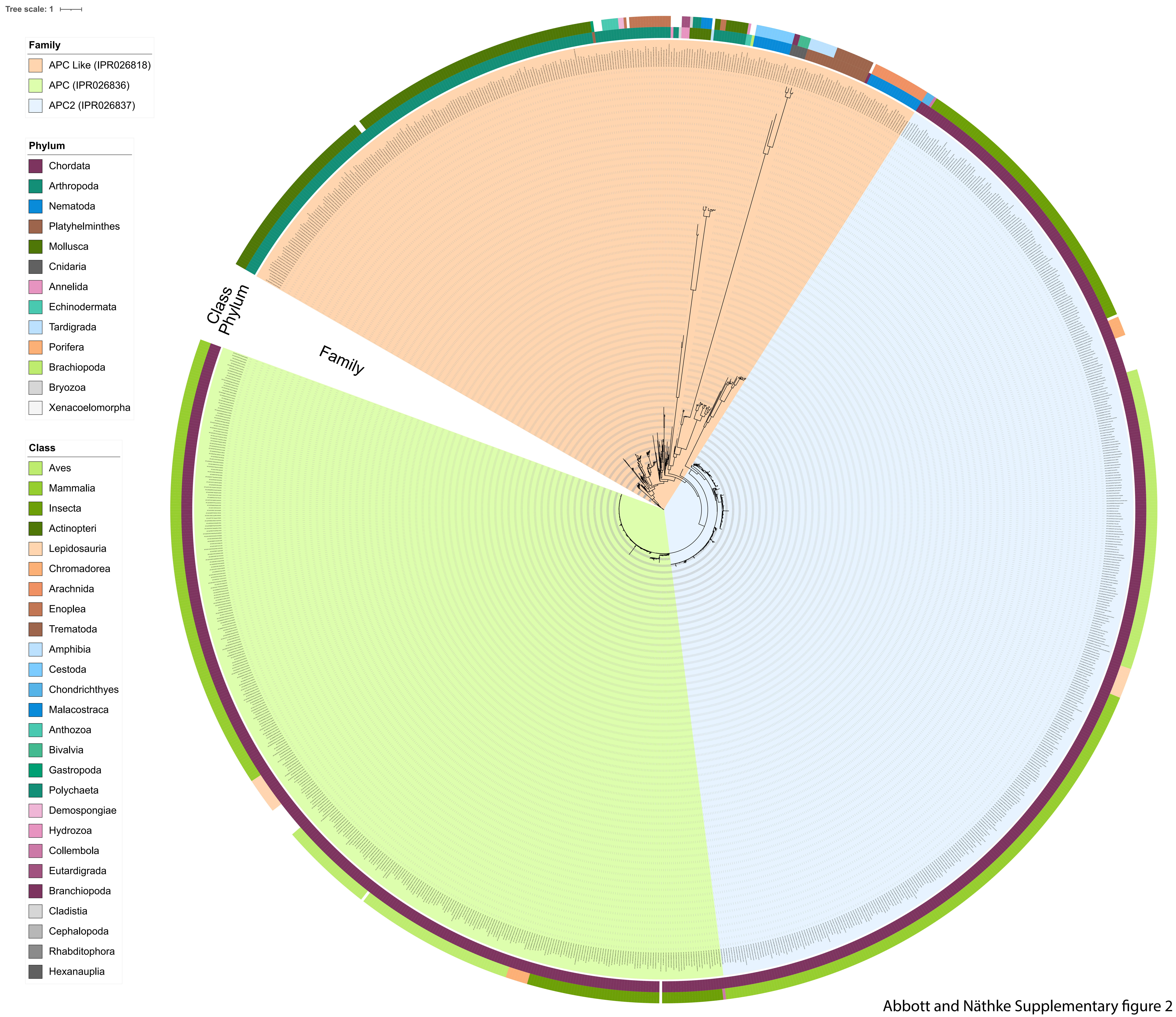


Abbott and Näthke Supplementary figure 2

### Supplemental figure 3

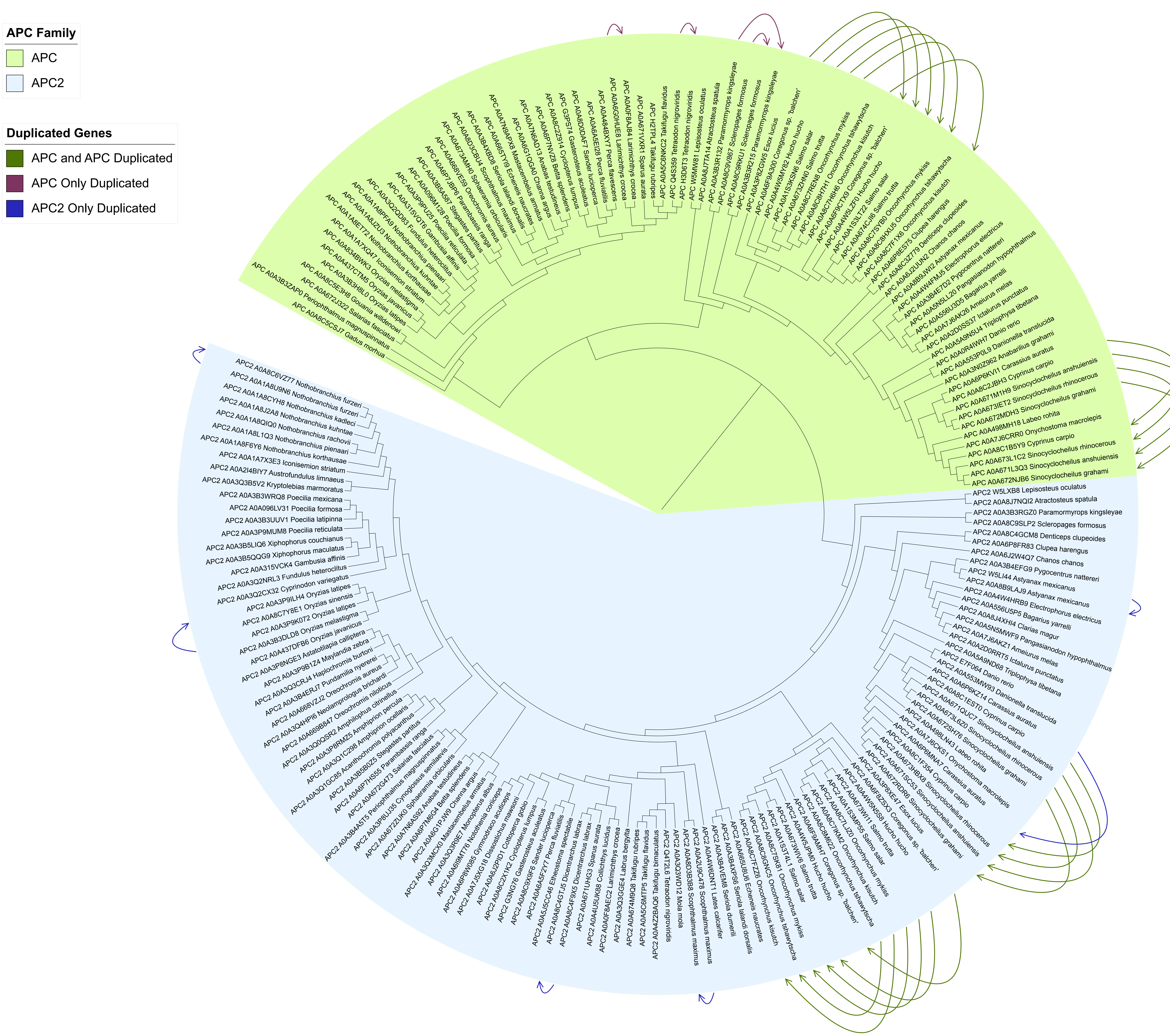


Abbott and Näthke Supplementary figure 3
